# Tuning selectivity of electrochemical sensors with polymer coatings

**DOI:** 10.64898/2026.02.16.706066

**Authors:** Ines C. Weber, Yann Zosso, Diego Uruchurtu Patino, Laura Rijns, Adrian L.M. Düsselberg, Zhenan Bao

## Abstract

Electrochemical sensors are promising for health monitoring due to their high repeatability and sensitivity, particularly when nanostructured. Yet, their translation into real applications is hindered by limited selectivity in the absence of specific binding receptors: many biomarkers exhibit similar oxidation potentials, producing overlapping voltammetric signals that impede molecular discrimination.

Here, we demonstrate that the oxidation potential of several small molecule biomarkers can be controlled through polymeric coatings, specifically poly(4-vinylpyridine), deposited onto glassy carbon electrodes. The polymer coating alters diffusion and adsorption characteristics, which ultimately lead to oxidation potential shifts of ascorbic acid and serotonin, enabling their separation of otherwise overlapping signals. These findings are supported by Chronocoulometry and

Fourier-transform infrared spectroscopy analysis that reveal changes in diffusion coefficient, adsorbed charge, and hydrogen bonding that are likely responsible for the altered sensor performance. These findings can be expanded to further polymers and biomarkers, including estradiol and melatonin. Finally, we demonstrate that the same selectivity trends persist on nanostructured, stretchable carbon-flower electrodes, where the high surface area further enhances sensitivity. Collectively, these findings reveal **polymer-controlled peak-potential tuning** as a powerful and broadly applicable route toward highly selective electrochemical sensors, enabling molecular discrimination in complex mixtures and opening new avenues for sensor-array-based detection.

## Introduction

Electrochemical sensors are among the most widely used sensor types. Their success stems from low cost, miniaturization potential, and operational simplicity: an applied potential drives redox reactions of target analytes, producing currents proportional to analyte concentration [1]. This process is repeatable, energy-efficient, and typically minimally disruptive to the sample. These characteristics make electrochemical sensors particularly attractive for point-of-care and wearable health monitoring [2, 3], enabling tracking of biomarkers for future personalized medicine. Promising results have been reported using non-invasive biofluids such as sweat [4, 5], saliva [6], tears, and breath [7].

Despite their potential, translation into real-world applications remains limited [8], with only a few technologies reaching clinical practice. **A major challenge is insufficient sensitivity and selectivity:** excellent ***sensitivity*** is required to detect trace-level concentrations of certain biomarkers in the (sub-)nanomolar (nM) range, while high ***selectivity*** is essential due to the complex composition of biofluids containing a plethora of biomarkers. Carbon-based materials have been widely studied in this context, and extensive efforts to improve sensitivity have focused on nanomaterial engineering [9–11], for example by tailoring morphology (carbon nanotubes [12], fuzzy graphene [13], carbon nanohorns [14]), or incorporating metal (oxide) dopants (i.e., Au [15, 16], Ag [16], Fe_3_O_4_[17, 18] and NiO [19]). These strategies enhance performance by increasing surface area, charge-transfer efficiency, and the density of electroactive sites, thereby promoting sensitivity. An alternative approach involves the use of conductive polymeric coatings to improve sensitivity, for example polyaniline [20], poly(3,4-ethylenedioxythiophene) [21], and polypyrrole [22].

While nano-engineering and conductive coatings have significantly improved sensitivity, selectivity often remains limited, as overlapping oxidation potentials of co-occurring biomarkers produce voltammetric signatures that cannot be reliably deconvoluted. Consequently, alternative strategies to enhance selectivity have been explored. Biosensors offer one such route through specific biorecognition [23], but often struggle with stability and repeatability because bound biomarkers do not readily detach. Data-driven approaches leveraging large datasets from single electrodes or arrays [24] have also been employed to improve selectivity. While these models enhance data interpretation and accuracy, they fail when oxidation peak overlap is substantial, particularly in scenarios where interferants are present at concentrations orders of magnitude higher than the target biomarker; for example, (sub-) nM serotonin (5-HT), melatonin (Mel), or estradiol (E2) in the presence of micromolar (µM) ascorbic acid (AA; vitamin C). Polymeric coatings have likewise been investigated, most notably the charged polymer Nafion®, which is used both for its antifouling properties and its ability to selectively attract or repel cations and anions, respectively [25, 26]. Overall, despite the merits of these approaches, we lack sensors that simultaneously meet stringent sensitivity and selectivity requirements. What is ultimately needed, and what motivates this work, is a strategy to deliberately control oxidation peak potentials, thereby enabling reliable deconvolution of overlapping electrochemical signals.

Here, we report the use of **neutral polymeric coatings** on carbon-based sensors to modulate the oxidation behavior of several biomarkers, including AA, 5-HT, E2, and Mel, thereby enabling improved selectivity. Specifically, we assess how poly(4-vinylpyridine) (P4VP) coated glassy carbon electrodes (GCEs) alter the peak potential and current of these biomarkers, demonstrating enhanced selectivity in mixtures. We further investigate the molecular interaction mechanism of the P4VP coating with the individual biomarkers through Chronocoulometry (CC) and Fourier-transform infrared spectroscopy (FTIR). We extend this concept to additional polymers such as poly(vinylidene fluoride) (PVDF), polystyrene (PS), and poly(2-vinylpyridine) (P2VP) to show the versatility of the approach. Finally, we demonstrate the applicability of this strategy using nanostructured carbon flowers, indicating that the polymer is the primary contributor to selectivity in carbon-based materials. These sensors exhibit enhanced sensitivity while preserving polymer-mediated selectivity. Our findings pave the way toward sensor arrays with distinct electrodes capable of selective multi-analyte detection.

## Results and discussion

### Concept

The rationale of our work is to modulate the oxidation potential and current through a polymeric coating. The polymer may interact with the biomarker through non-covalent interactions, such as van der Waals forces and hydrogen bonding, thereby modulating mass transport and the local electronic environment near the working electrode. On the one hand, reduced mass transport limits the concentration of molecules reaching the electrode surface and thus reduces the measured current. On the other hand, we hypothesize that polymer-biomarker interactions alter the electron distribution of the biomarker and its adsorption at the electrode interface, resulting in shifts in oxidation potentials [27].

This concept is schematically illustrated in **Fig. 1**. In the absence of a coating, biomarkers freely diffuse to the electrode and undergo oxidation at partially overlapping potentials, hindering deconvolution in mixtures (**Fig. 1a**). In the presence of a polymeric layer, mass transport and hence diffusion coefficients as well as the local electronic environment at the electrode interface are altered, affecting peak current and separating oxidation potentials, therefore, improving selectivity (**Fig. 1b**).

**Fig. 1.**
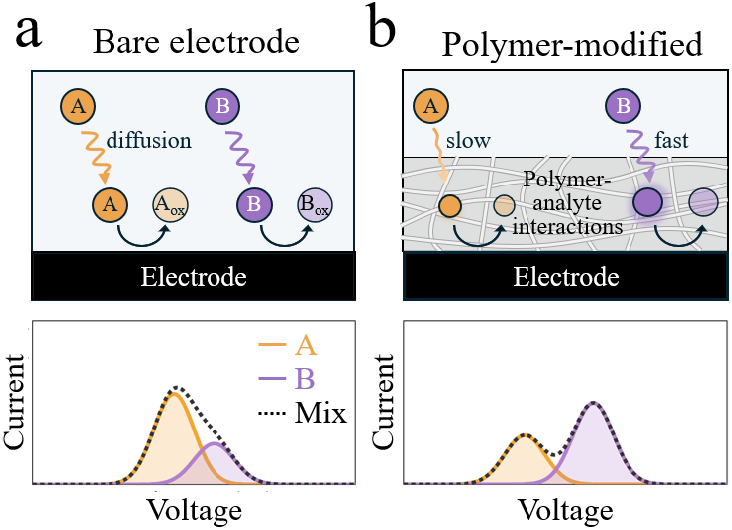
Concept. **(a)** On a bare electrode, two biomarkers diffuse to the surface and are oxidized at similar potentials, producing overlapping voltammetric signals. **(b)** A polymer coating modulates diffusion and the local electronic environment of each biomarker, resulting in selective detection.

### Modulating oxidation potentials with P4VP

P4VP was chosen as a polymer coating because its pyridinic nitrogen can interact with a wide range of molecules, acting as a strong Lewis base and an efficient hydrogen-bond acceptor. AA and 5-HT were chosen as model biomarkers because of their clinical relevance [28, 29] and differing physiological concentrations. Specifically, AA occurs in the µM range in saliva after fasting [30] and fluctuates significantly in sweat upon supplementation [31]; whereas peripheral 5-HT levels are in the low nM range in saliva [32] and hardly assessed at all in sweat. This concentration disparity makes selective detection of 5-HT particularly challenging.

**Figure 2a** shows normalized square-wave voltammograms obtained with an uncoated GCE exposed to AA and 5-HT *separately* (i.e., sequential measurements; raw data **Fig. S1a**). AA is detected at 0.08 ± 0.01 V, while 5-HT exhibits a characteristic two-peak response at 0.30 and 0.39 ± 0.01 V previously attributed to sequential oxidation to a carbocation and then to a quinone imine [33]. The AA and 5-HT signals overlap in the range of 0.2 to 0.4 V. Consequently, when measured *in mixtures* containing 500 µM AA and increasing 5-HT concentrations (500 – 3000 nM; **Fig. 2b**, raw data **Fig. S1b**), the 5-HT signal becomes obscured by the broad AA shoulder due to both peak overlap and the large concentration disparity. Note that these concentration ranges were deliberately chosen to challenge the sensor, as in biological environments ascorbic acid can be present at concentrations orders of magnitude higher than serotonin at levels selected here.

**Fig. 2.**
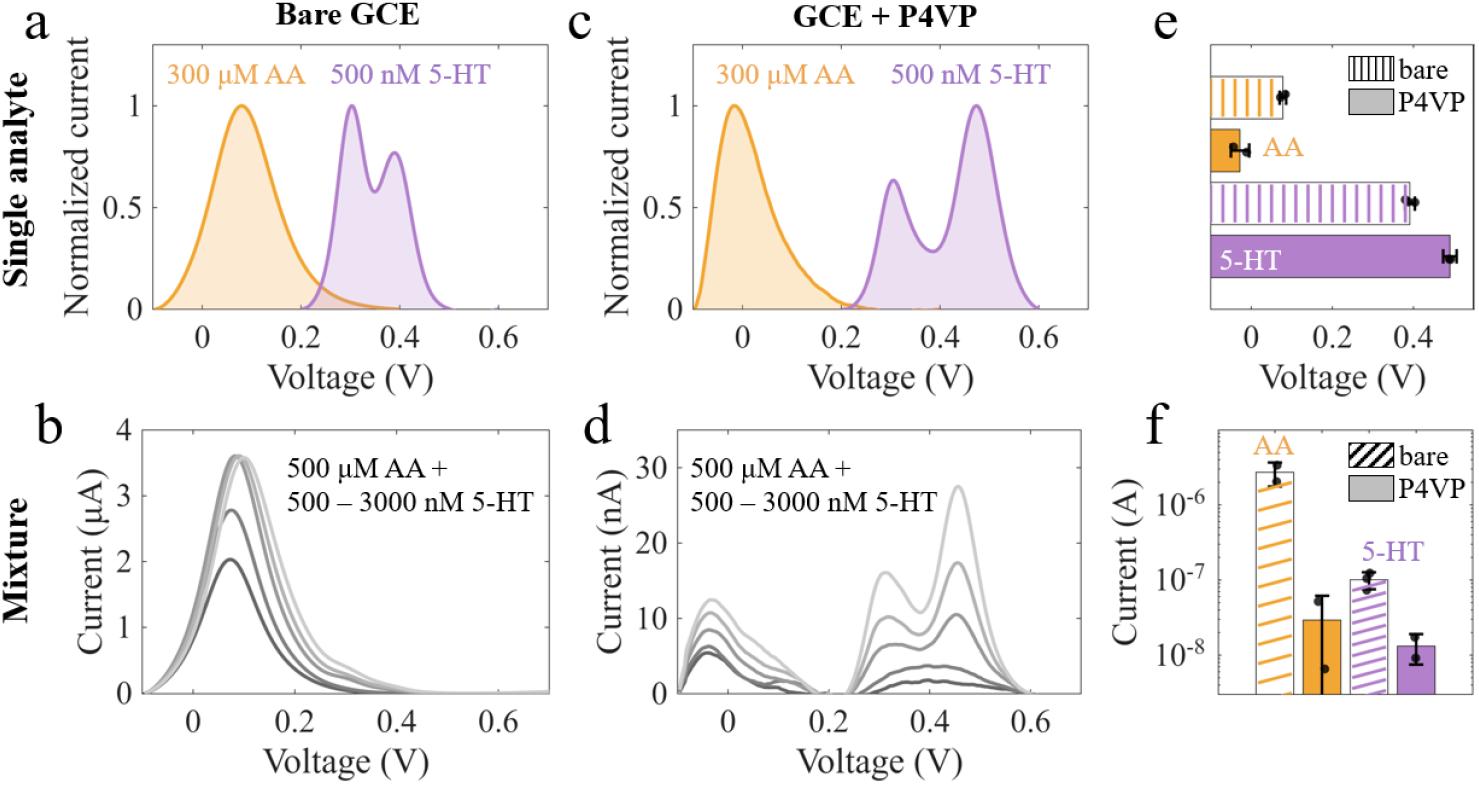
Polymer shifts oxidation peak potential. **(a)** Single-analyte SWV scans of 300 μM AA or 500 nM 5-HT on bare GCE plotted together. **(b)** SWV of AA/5-HT mixtures containing 500 μM AA with increasing 5-HT concentrations (500 − 3000 nM). **(c,d)** Analogous SWV scans for the P4VP-modified electrode. **(e,f)** Comparison of peak potentials and peak currents for AA and 5-HT in the absence and presence of the P4VP.

Introducing a P4VP coating dramatically alters the system’s electrochemical response. The oxidation potential of AA shifts negatively to −0.03 ± 0.02 V, while that of 5-HT shifts positively to 0.31 and 0.49 ± 0.01 V, thereby increasing the peak separation and enabling clear discrimination (**Fig. 2c**, raw data **Fig. S1c**). It is worth noting that these potential shifts appear independent of the P4VP coating thickness (**Fig. S2a**), suggesting that interactions between P4VP and the biomarkers at the electrode surface determine the peak positions rather than the coating thickness. This is advantageous because it minimizes electrode-to-electrode variability and may allow the use of thin polymer coatings that modulate peak potential without significantly hindering diffusion, thereby preserving sensitivity. The relevance of these peak shifts is reflected when measuring the same biomarkers in mixture (again 500 – 3000 nM 5-HT, **Fig. 2d**, raw data **Fig. S1d**), where increasing 5-HT concentrations remain clearly visible despite the presence of high AA concentrations. Notably, particularly AA but also 5-HT currents decrease substantially upon P4VP modification (from 2.7 microamperes (μA) to 29 nanoamperes (nA) for AA), and similar currents are observed with varying P4VP thickness (**Fig. S2b**). The current reduction is also consistent with the trends observed in electrochemical impedance spectroscopy (EIS) measurements (**Fig. S2c**).

In summary, the P4VP modification leads to a noticeable AA shift towards lower potentials, while 5-HT shifts towards higher potentials (**Fig. 2e**). Simultaneously, the current decreases for AA by two orders of magnitude, while it decreases by one order of magnitude for 5-HT (**Fig. 2f**). Hence, the **polymeric coating modulates both oxidation potential and current amplitude**, two powerful handles for improving selectivity, as shown on the example of AA and 5-HT.

### Mechanistic considerations

To investigate how P4VP modifies interfacial interactions at the working electrode, we employed double potential-step chronocoulometry (CC). CC is a well-established technique to evaluate apparent diffusion coefficients of redox-active molecules as well as their adsorption on the working electrode [34].

The CC response of bare and P4VP-coated GCEs in the presence of AA (2000 μM) and 5-HT (200 μM) is shown in **Fig. 3a**. In both cases, the total charge is reduced after coating the electrode. This decrease likely reflects hindered access of the biomarkers to the electrode surface, caused by slower transport through the polymer layer. These observations are consistent with the increased semicircle observed in EIS (**Fig. S2c**) and the reduced voltammetric currents in Figure 2.

**Fig. 3.**
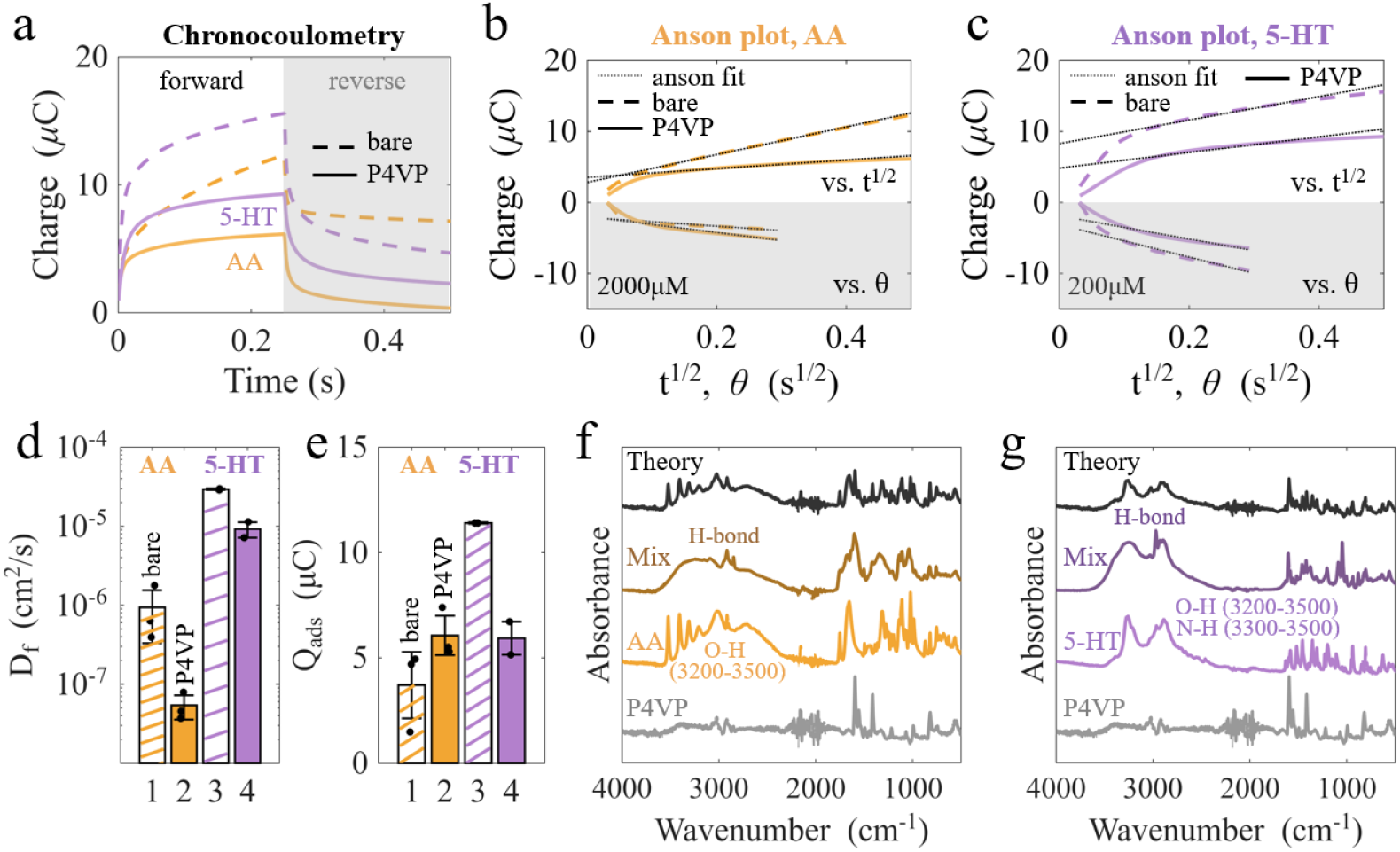
Mechanistic considerations. **(a)** Chronocoulometry of 200 μM 5-HT and 2000 μM AA on bare GCE (dashed) and P4VP-coated GCE (solid line). **(b)** Corresponding Anson plots for AA. **(c)** Corresponding Anson plot for 5-HT. **(d)** Derived diffusion coefficients (D_f_). **(e)** Derived adsorbed charge (Q_ads_). **(f)** FTIR measurements of P4VP, ascorbic acid, their mixture, and theoretical self-sorted spectra. **(g)** Analogous for 5-HT.

The corresponding Anson plots [35] for AA and 5-HT are shown in **Fig. 3b,c**. The slopes of the linear regions (dotted lines) yield the apparent diffusion coefficients. For both biomarkers, the P4VP coating lowers the slope, indicating a reduced diffusion coefficient (**Fig. 3d**). Notably, AA exhibits a pronounced decrease in *D*_*f*_ from 9.4 × 10^-7^ cm^2^/s to 6.08 × 10^-8^ cm^2^/s upon P4VP modification, whereas 5-HT shows a more moderate reduction, from 2.9 × 10^-5^ cm^2^/s to 6.6 × 10^-6^ cm^2^/s (**Fig. 3e**). The differences in *D*_*f*_ between AA and 5-HT at the bare electrode may arise from dissolution and mass transport of these biomarkers in the P4VP and could be influenced by concentration difference (2000 μM vs 200 μM, respectively); the chosen concentration ranges were selected because 5-HT typically produces stronger signals than AA. However, the stronger reduction of *D*_*f*_ for AA compared to 5-HT again is consistent with the greater current suppression observed for AA relative to 5-HT (**Fig. 2f**).

The adsorbed charge *Q*_*ads*_, obtained from the intercept at *t = 0*, further highlights differences in polymer-biomarker interactions. For AA, *Q*_*ads*_ increases in the presence of P4VP, whereas for 5-HT it decreases. Thus, P4VP enhances AA adsorption but reduces it for 5-HT. The increased adsorption of AA is consistent with prior reports of pyridine-based polymers promoting hydrogen-bond-stabilized adsorption of enediol species [27]. Notably, changes in *Q*_*ads*_ correlate with the observed shifts in oxidation potential: increased adsorption tends to lower the oxidation potential, whereas reduced adsorption shifts it to more positive values.

While the exact relationship between adsorbed charge and oxidation potential depends on the kinetic regime, such trends are expected because adsorbed charge can modulate heterogeneous electron-transfer kinetics in irreversible systems [34]; in fact, a previous study has shown a correlation between adsorbed charge and electron-transfer rates using CC [35].

Beyond CC, we employed FTIR to study the chemical interactions of P4VP with the biomarkers in 1:1 molar mixture (**Fig. 3f,g**). Prior to FTIR analysis, all samples were thoroughly dried in a vacuum oven (40-50 °C, 1 h) to ensure removal of solvents and water. The spectra of the pure biomarkers are in good agreement with literature [36], where AA exhibits its characteristic O–H stretching bands at 3215–3520 cm^−1^[37], and 5-HT shows overlapping O–H and N–H stretching bands in the ∼3200–3500 cm^−1^region [38].

In contrast, the combination of P4VP with biomarker mixtures display clear spectral changes. Both AA–P4VP and 5-HT–P4VP mixtures show pronounced broadening and a redshift of the O–H/N–H stretching region (3000-3500 cm^−1^), reflecting the formation of hydrogen bonds. These interactions are chemically plausible: the pyridinic nitrogen in P4VP is a strong hydrogen-bond acceptor, and both biomarkers provide donor groups (i.e., O– H in AA and both O–H and N–H in 5-HT) enabling interactions of the form O–H⋯N=C or N–H⋯N=C.

To confirm that the polymer and biomarker are indeed interacting, rather than co-existing in a self-sorted state, we compared the experimental spectra with “theoretical” spectra generated by molar averaging of the pure components [39] by simply overlaying the separately measured spectra (in theory non-interacting). They deviate substantially, supporting the presence of P4VP-biomarker interactions and not a mere non-interacting state.

The formation of hydrogen bonds indicates that the polymer alters the local interfacial environment. For 5-HT, such hydrogen bonding likely withdraws electron density from the indole unit, making oxidation less favorable and requiring higher potentials. The AA, however, shifts towards lower potentials despite also forming hydrogen bonds. This may reflect stronger interfacial accumulation or preferential adsorption within the P4VP layer, which can lower the apparent oxidation potential under mixed kinetic–diffusion control.

In summary, CC and FTIR together provide initial insights into the mechanisms underlying the voltammetric changes. CC data reveals stronger reduction in AA diffusion compared to 5-HT, consistent with the observed sensing behavior, while adsorbed charges correlate with potential shifts. FTIR confirms hydrogen bonding between the polymer and the biomarkers, which alters the local interfacial environment. Further mechanistic studies could resolve the individual contributions of these factors.

### Expanding to other polymers and biomarkers

A key feature of this work is the versatility of tuning polymer-biomarker interactions. The simplicity of the approach enables extension beyond AA and 5-HT to other biomarkers, and beyond P4VP to other polymers, yielding a flexible toolbox of polymer-biomarker combinations with tunable selectivity and sensitivity.

The evaluated peak potentials are shown on the example of AA in **Fig. 4a** with various polymer modifications (full SWV scans for all biomarkers in **Fig. S4**). Both pyridine-containing polymers (i.e., P4VP and P2VP) exhibit a negative peak shift, reaching potentials as low as −0.04 V. This further verifies the polymer-biomarker interactions discussed earlier for P4VP as similar interactions are expected for P2VP. Polystyrene (PS) shows negligible change in peak potential, consistent with expected minimal interaction between the styrene ring and AA. In contrast, poly(vinylidene fluoride) (PVDF) shifts the oxidation peak to significantly higher potentials. While the origin of this effect requires further investigation, several factors may contribute, including the hydrophobic nature of PVDF, which may slow down the diffusion of the hydrophilic AA.

**Fig. 4.**
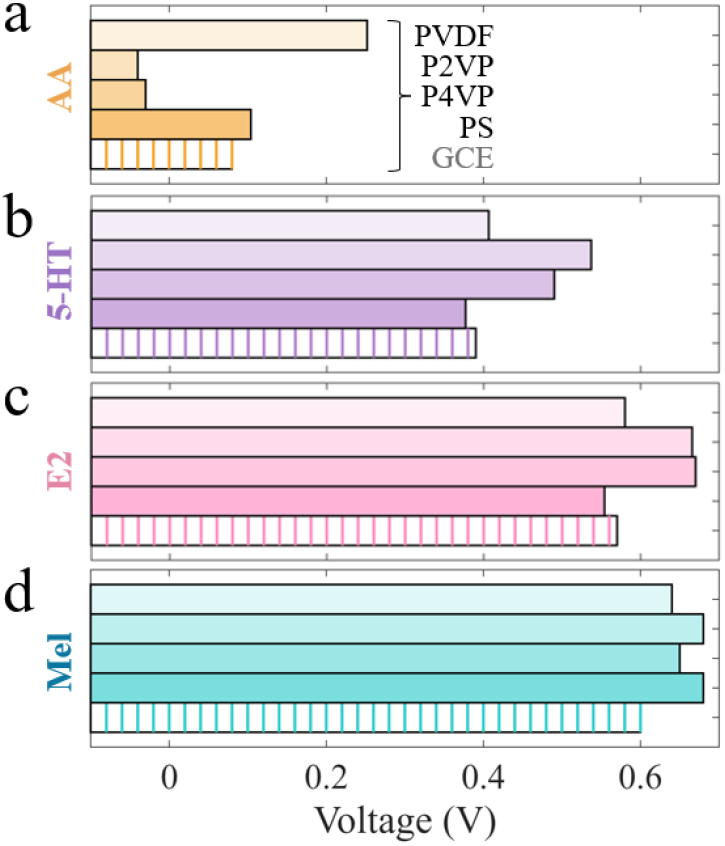
Peak potential shifts. **(a)** Peak locations for PVDF, P2VP, P4VP, PS, and bare GCE for AA. **(b)** Analogous for 5-HT. **(c)** Analogous for E2. **(d)** Analogous for Mel.

In case of 5-HT, both pyridine-containing polymers shift the oxidation potential to higher values (**Fig. 4b**). Again, PS shows minimal impact on peak potential. Finally, PVDF-coatings lead to a slight negative peak shift, which is however less pronounced compared to AA. Note that with all polymer coatings, two peaks are observed for 5-HT oxidation, and the second peak is used to evaluated the oxidation potential.

Furthermore, we investigated melatonin (Mel) and estradiol (E2), two biologically relevant biomarkers due to their roles in sleep regulation [40] and female health [41], respectively. Their electrochemical behavior is comparable to that of 5-HT: the largest positive shifts are observed for pyridine-containing polymers, whereas PS and PVDF induce only slight positive shifts or negligible effects (**Fig. 4c,d**). Notably, both biomarkers are capable of hydrogen-bonding with P4VP (**Fig. S4**), which may reduce the electron density of the biomarkers and contribute to the observed positive potential shifts. It is also noteworthy that PVDF has a minimal impact on E2 detection, consistent with E2 being the most hydrophobic biomarker among the four biomarkers tested. Overall, these results demonstrate neutral polymer modifications as a versatile strategy for tuning peak potential and selectivity. Future work will expand this toolbox using polymers with diverse functional groups.

### Universal: from GCE to carbon flowers

GCEs offer a well-established electrode platform for studying basic electrochemical effects as discussed earlier. However, due to their bulky and rigid nature, as well as the flat surface giving limited sensitivity, they are unsuitable for wearable sensors in health applications. Hence, we expanded polymeric coatings beyond GCE and incorporated them together with carbon flower (CF) electrodes. CFs exhibit excellent sensing performance [42] due to their unique morphology, high conductivity, tunable size [43], and accessible surface area, as shown in the SEM image in **Fig. 5a**. In addition, these carbon flowers can be readily processed by spray coating techniques (**Fig. 5b**) onto stretchable substrates (**Fig. 5c**), making them well suited for skin-mounted applications. Polymer modifications can be directly incorporated into the spray-coating solution, where they also serve as binders between the carbon flowers, enabling one-step fabrication of sensing electrodes. A drawback of this co-deposition approach is the limited control over polymer distribution, which can lead to inhomogeneous coatings, as evidenced by scanning electron microscopy (SEM, **Fig. 5d** and **Fig. S5**).

**Fig. 5.**
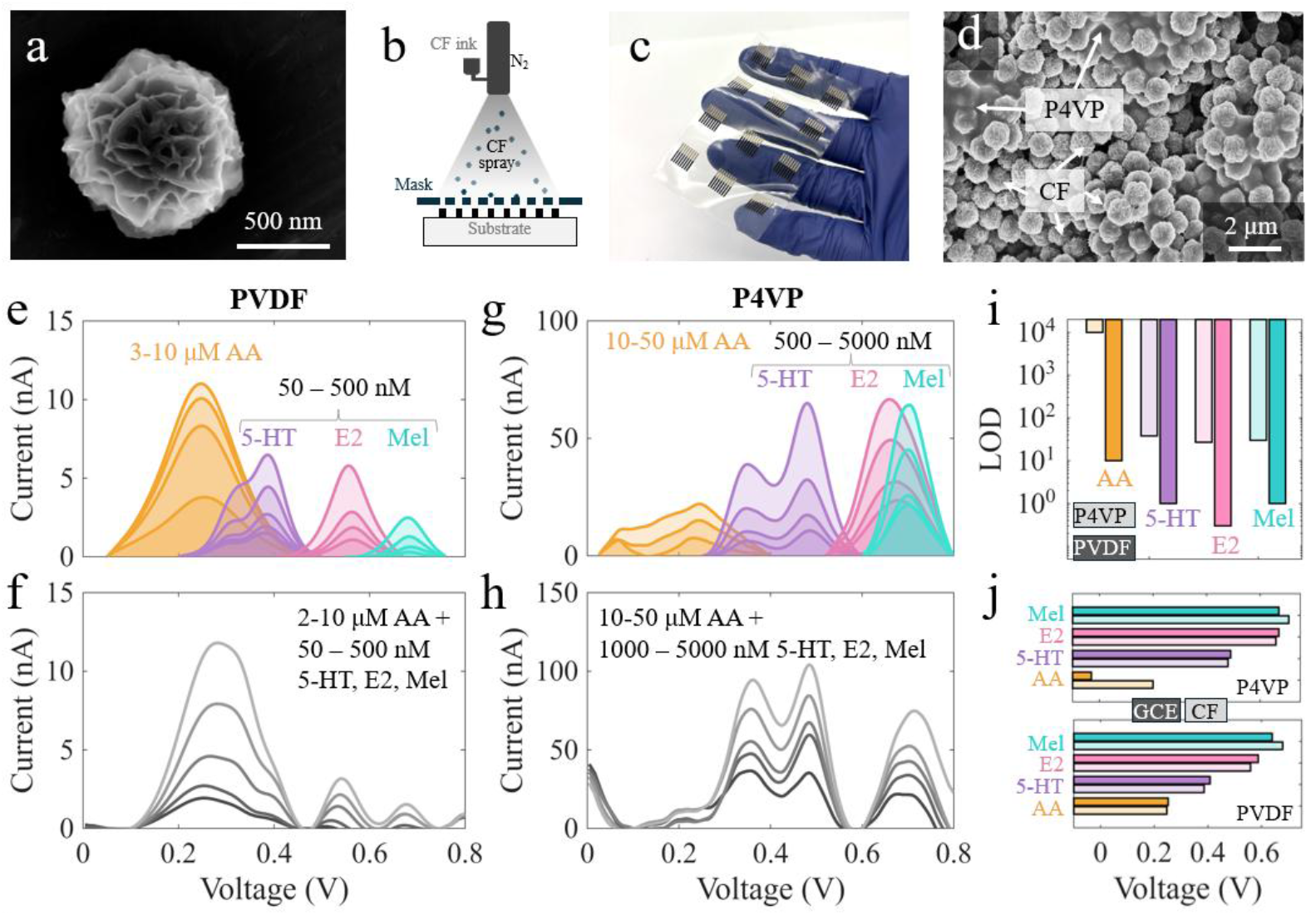
Stretchable carbon flower (CF) sensors. **(a)** SEM image of a single CF. **(b)** Schematic of CF sensor fabrication via spray coating. **(c)** Photograph of flexible sensing electrodes. **(d)** SEM image of a P4VP-CF electrode. **(e)** Single-analyte voltammograms plotted together obtained with a PVDF-CF electrode for AA, 5-HT, E2, and Mel. **(f)** Corresponding mixture voltammograms. **(g)** Single-analyte voltammograms plotted together obtained with a P4VP-CF electrode for A A, 5-HT, E2, and Mel. **(h)** Corresponding mixture voltammograms. **(i)** Limits of detection (LOD) of PVDF-CF and P4VP-CF. **(j)** Oxidation potentials for P4VP or PVDF on GCE and CF.

Despite inhomogeneous polymer deposition, polymer-induced changes to sensing performance are retained: **Fig. 5e** shows CF sensor responses with PVDF for single biomarker exposures: 3–10 µM AA and 50–500 nM 5-HT, E2, and Mel (raw data **Fig. S6a-d**). While all biomarkers are detectable on PVDF-CF electrodes, AA and 5-HT signals significantly overlap, as seen in the mixture measurements (**Fig. 5f**, raw data **Fig. S6e**), where the two peaks merge.

In contrast, P4VP-CF electrodes reduce the AA response while shifting 5-HT, E2, and Mel towards larger potentials (**Fig. 5g**, raw data **Fig. S6f-i**). Here, mixture measurements allow clear distinction between AA and 5-HT, while E2 and Mel merge (**Fig. 5h**, raw data **Fig. S6j**). Note that higher biomarker concentrations were needed for P4VP-CF due to generally noisier signals. This is reflected in **Fig. 5i**, showing better (i.e., lower) limits of detection (LOD) for PVDF compared to P4VP. The difference in LOD should be attributed to both the polymer modification (i.e., reduced diffusion) and the fabrication process itself, as the polymer also acts as binder; further investigations are needed to optimize the co-deposition of carbon flowers and polymers to enhance signal sensitivity. However, both PVDF and P4VP feature good sensitivities in the lower nM range, sufficient for most clinical applications.

It is important to note that the observed oxidation potentials are in excellent agreement with GCE electrodes (**Fig. 5j**). A small exception is presented by AA on P4VP-CF electrodes, where however the signal is so small that precise peak potential determination remains challenging. Overall, the simple addition of polymer binders, even when distributed inhomogeneously, markedly improves sensor selectivity while retaining high sensitivity and flexibility. With more controlled polymer deposition, these coatings could reliably tune peak potentials without compromising (and potentially even enhancing) sensitivity, for example by increasing adsorbed charge. This strategy provides a foundation for fabricating selective sensor arrays by combining electrodes modified with different polymers.

## Conclusion

Carbon-based electrochemical sensors often suffer from limited selectivity due to overlapping biomarker signals. Here, we demonstrate that neutral polymer coatings provide an effective strategy to modulate oxidation potentials and thereby enhance selectivity. Specifically, P4VP reduced the oxidation potential for AA, while 5-HT oxidation is shifted towards higher voltages, enabling clear discrimination between these two otherwise overlapping biomarkers.

Mechanistic investigations using CC and FTIR spectroscopy reveal that P4VP influences both the apparent diffusion coefficient and the adsorbed charge. Specifically, P4VP enhances the adsorbed charge of AA while reducing that of 5-HT, correlating with the observed peak shifts. FTIR measurements further indicate hydrogen bonding between the biomarkers and P4VP, suggesting that such non-covalent interactions play a key role in modulating oxidation potentials.

This concept extends beyond P4VP to other polymers, such as PVDF, P2VP, and PS, and to additional biologically relevant biomarkers including E2 and Mel. Importantly, the approach is transferable across carbon materials, as demonstrated on carbon flower electrodes, where similar peak shifts and selective detection of four biomarkers (AA, 5-HT, E2, and Mel) are achieved with high sensitivity.

In summary, introducing polymer coatings onto sensors provides a powerful and versatile means to **enhance the selectivity** of electrochemical sensors. This approach is particularly attractive for real fluids such as sweat and saliva, where biomarkers co-occur at widely varying concentrations. Consequently, this strategy offers a promising pathway toward the development of selective sensor arrays and wearable health-monitoring devices.

## Experimental

### GCE electrode preparation

GCEs (2 mm diameter) were obtained from CH Instruments Inc. Prior to modification, the electrodes were mechanically polished using alumina slurries (0.3 µm and 0.05 µm) from the CH Instruments polishing kit. After polishing, the electrodes were thoroughly rinsed with deionized water to remove any residual alumina particles.

Solutions of P4VP at concentrations of 2, 5, and 10 g/L were prepared in chloroform. Each GCE was then coated by drop-casting 16 µL of the respective polymer solution onto the electrode surface and allowing the solvent to evaporate. Additional polymer-modified GCEs, including P2VP, PS, and PVDF, were prepared using the same procedure, employing polymer solutions at a concentration of 5 g/L.

### Electrochemical testing

Electrochemical experiments were carried out using a CH Instruments 760E potentiostat equipped with a CHI200 picoamp booster and Faraday cage. A standard three-electrode setup was used, consisting of either the GCE electrodes or the carbon flower electrodes, an Ag/AgCl (3 M KCl) reference electrode, and a platinum wire counter electrode. Measurements were performed in nitrogen-degassed 1× PBS (pH 7.2) at room temperature.

L-ascorbic acid (≥98%), serotonin hydrochloride (≥98%), melatonin (≥98%), and 17β-estradiol (>97%) were prepared immediately before use (in PBS for the first two; ethanol for the latter two). Twenty background scans in pure PBS were recorded to stabilize the baseline. Biomarker additions were made sequentially, with gentle mixing achieved by brief nitrogen bubbling.

SWV was employed for biomarker detection due to its high sensitivity. For GCE-based electrodes, scans were recorded from 0.1 V to 0.9 V, and for CF-based electrodes from 0 V to 0.8 V. All measurements used a potential step of 0.004 V, an amplitude of 0.025 V, and a frequency of 5 Hz. Each SWV scan was performed in triplicate, with a 60 s quiet time between scans and following each concentration increase.

EIS measurements were performed at the open-circuit potential using a frequency range of 100 kHz to 0.1 Hz and an alternating current perturbation amplitude of 10 mV.

Double potential-step CC was performed to determine the apparent forward diffusion coefficient (*D*_*f*_) and the adsorbed charge (*Q*_*ads*_). A blank scan in PBS (without biomarker) was collected prior to each experiment. The pulse width was set to 250 ms with a sampling interval of 1 ms. The applied potential steps were biomarker-dependent: for AA and 5-HT, Ei = −0.2 V and Ef = 0.6 V were used; for E2 and Mel, Ei = 0.3 V and Ef = 0.9 V were applied. The apparent forward diffusion coefficient was calculated from the slope of the forward-step charge–time response:

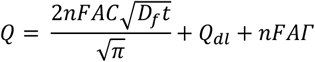

Where *Q* is the total charge in Coulomb that has passed at time t, *n* is the number of electrons transferred, *F* is the Faraday constant, *A* is the electrode area, *C* is the biomarker concentration, *D*_*f*_ is the apparent forward diffusion coefficient, *Q*_*dl*_ is the double-layer charge and Γ is the amount of adsorbed reactant. The adsorbed charge (*Q*_*ads*_) was obtained from the difference in the fitted intercepts of the forward (*Q*_*int,f*_) and reverse Anson plot (*Q*_*int,r*_):

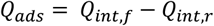

The reverse charge is obtained by:

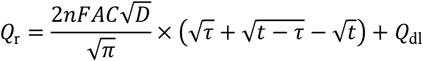

where τis the switching time between forward and reverse potential steps.

### Carbon flower sensors

Carbon flower (CF) materials were synthesized following our previous procedures [44]. Briefly, acrylonitrile and acetone (1:1 v/v) with AIBN (1 mg mL^−1^) were reacted at 70 °C for 4 h in sealed vials to form polymeric flowers, which were then washed with methanol, dried, oxidatively stabilized at 230 °C for 4 h, and carbonized at 1500 °C for 2 h.

CF ink was prepared by dispersing 45 mg of CF material and 5 mg PVDF in 100 mL chloroform, followed by overnight bath sonication and probe sonication (20 min, 40% amplitude, 2 s on/off). The mixture was then centrifuged (1000 rpm, 1 min), and the resulting supernatant was collected for subsequent electrode fabrication.

Flexible sensors were fabricated by drop-casting SEBS (150 mg mL^−1^ in cyclohexane) onto glass slides and drying overnight. A laser-cut Kapton mask defining eight electrodes (10.5 × 0.5 mm) was applied, and stretchable silver paste was blade-coated and cured at 100 °C for 2 h. After mask removal, a precision metal mask was aligned over the silver traces and fixated with a magnet, and 7 mL of CF ink was spray-coated at 80 °C. The mask was removed, and devices were encapsulated either by spin-coating SEBS (80 mg mL^−1^) or applying Kapton tape. Electrical connections were made using conductive z-axis tape.

### Material characterization

An FEI Magellan 400 XHR scanning electron microscope was used for SEM imaging (100 pA, 5 kV). FTIR spectra were collected in the solid state using an ATR accessory over 4000– 500 cm^−1^ with ∼0.1 cm^−1^resolution, averaging 16 scans per sample, on a Cary 6000i Spectrophotometer.

For FTIR sample preparation, P4VP was mixed at a 1:1 molar ratio with each biomarker. A 10 g/L P4VP solution in chloroform was used, and biomarkers were either pre-dissolved in water–ethanol mixtures or added directly to chloroform (Mel) to ensure fully transparent, homogeneous solutions. Solvents were evaporated in a vacuum oven (40–50 °C, 1 h, cap open), and the resulting dry powder was placed directly on the ATR crystal for measurement.

## Data analysis

SWV data were processed by subtracting the background response obtained in blank PBS under identical conditions. After background correction, a linear baseline between the start and end of each peak was removed to isolate the biomarker signal. Data were smoothed using a 20-point moving average (“movmean” in MATLAB). Analytical sensitivity was determined from the slope of peak current versus concentration, and the limit of detection (LOD) was calculated as LOD = 3σ/S, where σ is the baseline noise and S is the analytical sensitivity. All data analysis was performed in MATLAB (2023), with replicate measurements shown as individual data points.

## Acknowledgements

This work was supported by a seed grant from the Stanford Wearable Electronics Initiative (eWEAR) and by postdoctoral fellowship funding from the Swiss National Science Foundation (SNSF) to I.C.W. (P500PT_214498), as well as from the Niels Stensen Fellowship and the Netherlands Organisation for Scientific Research (NWO) through the Rubicon Fellowship (019.233EN.013) to L.R. Z.B. is a Chan Zuckerberg Biohub San Francisco investigator and an Arc Institute innovation investigator. Portions of this research were carried out at the Stanford Nano Shared Facilities (SNSF), supported by the National Science Foundation under award ECCS-2026822. We thank F. Stellacci, S. Lacour, M. Cargnello, K. Parkatzidis, K.-J. Hsu, and A. Mahmud for valuable scientific discussions.

## Data availability

The data that support the findings of this study are available from the corresponding author upon request.

