## Supplementary Information for "Tuning selectivity of electrochemical sensors with polymer coatings"

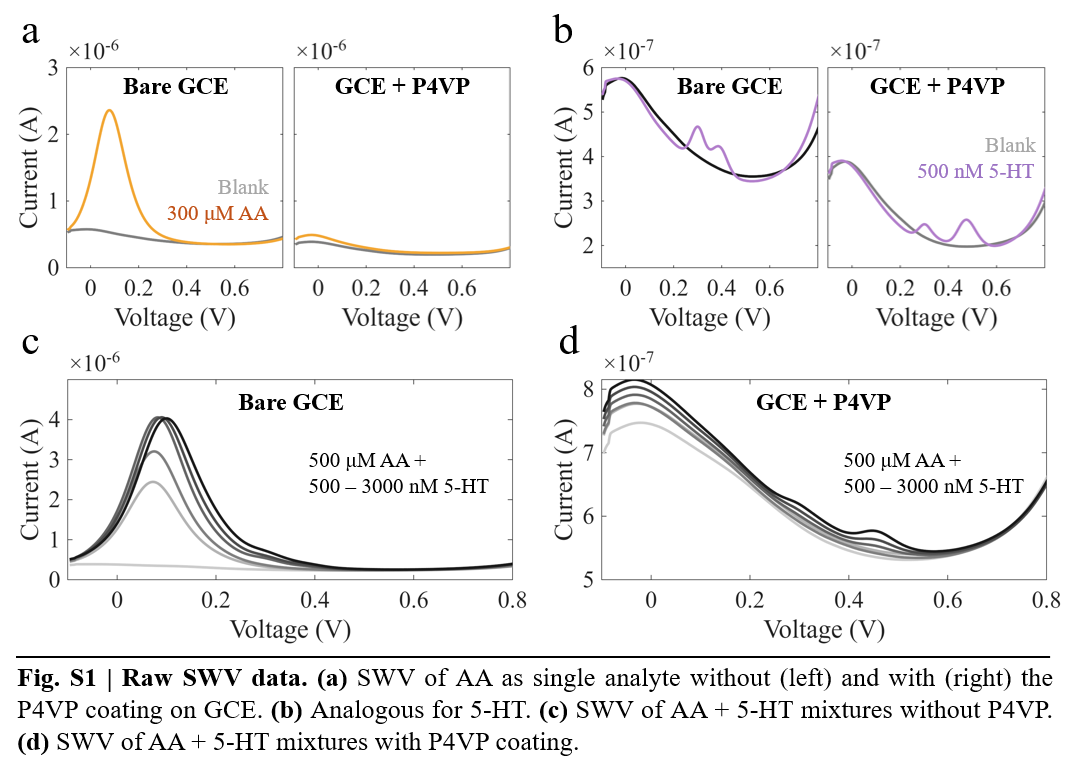

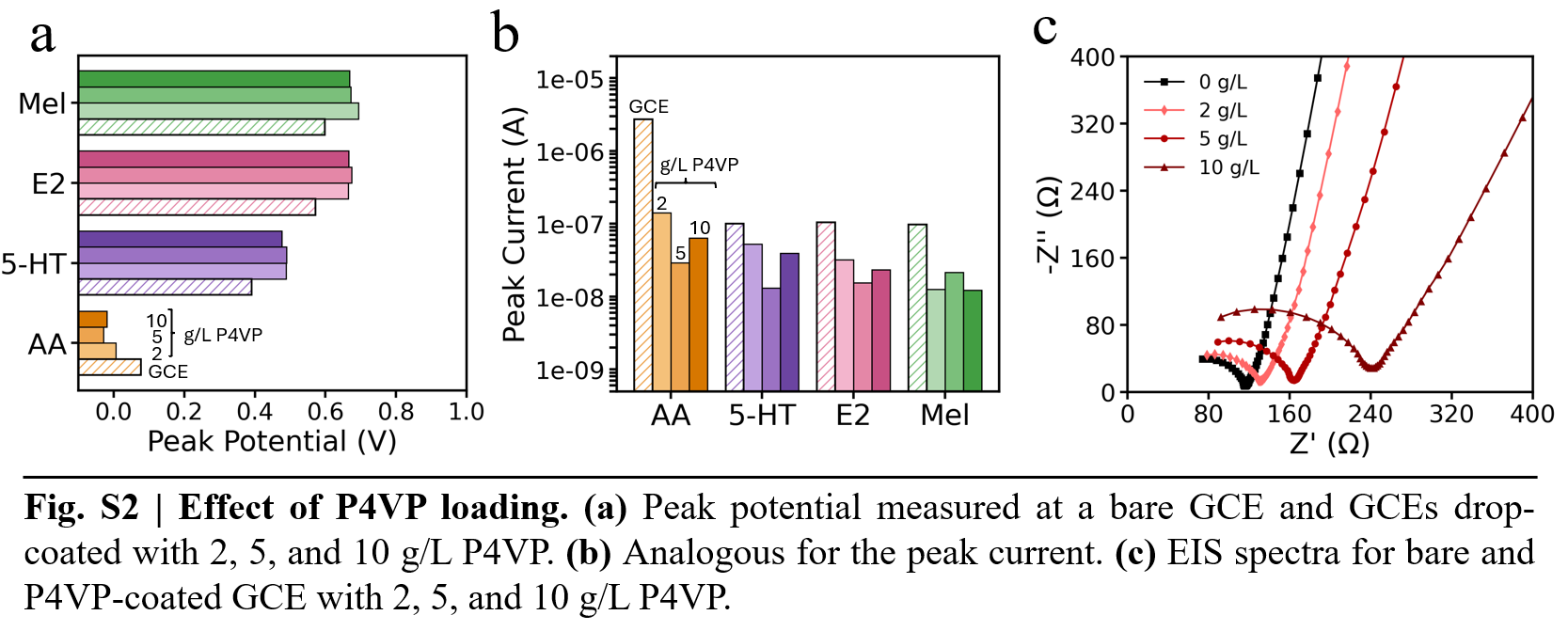

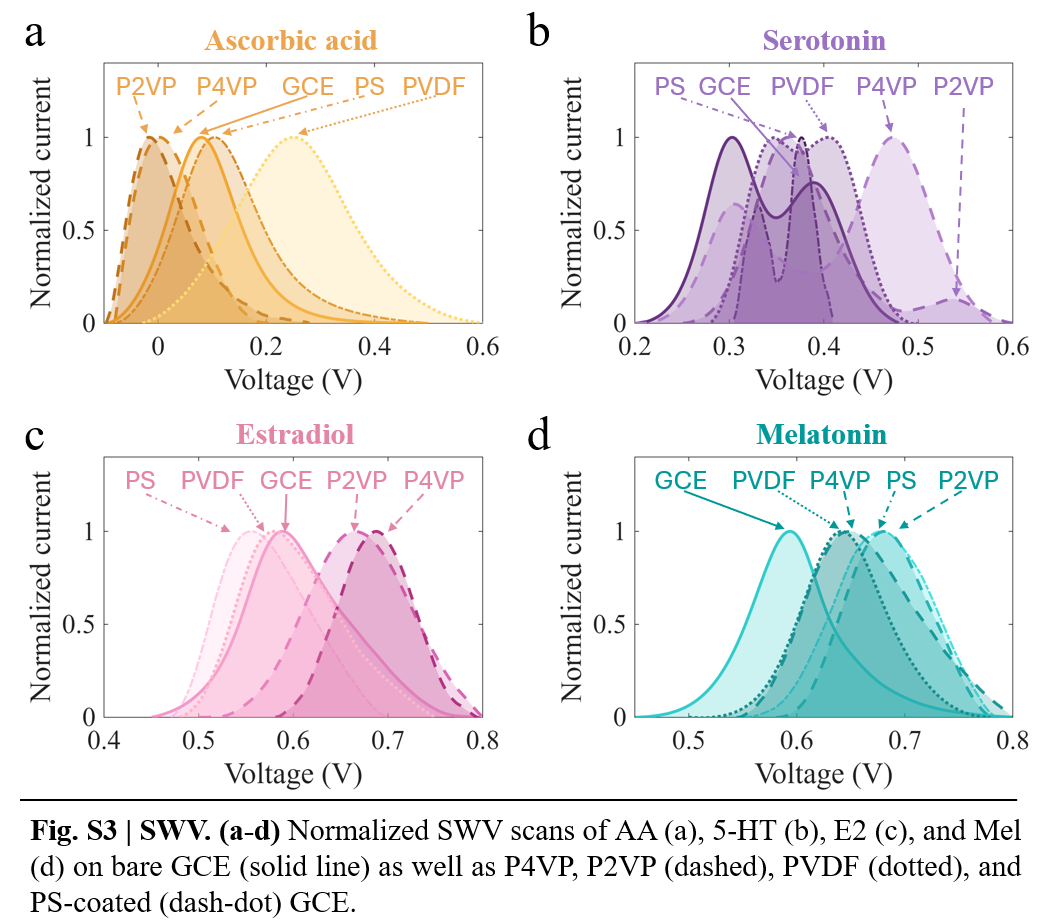

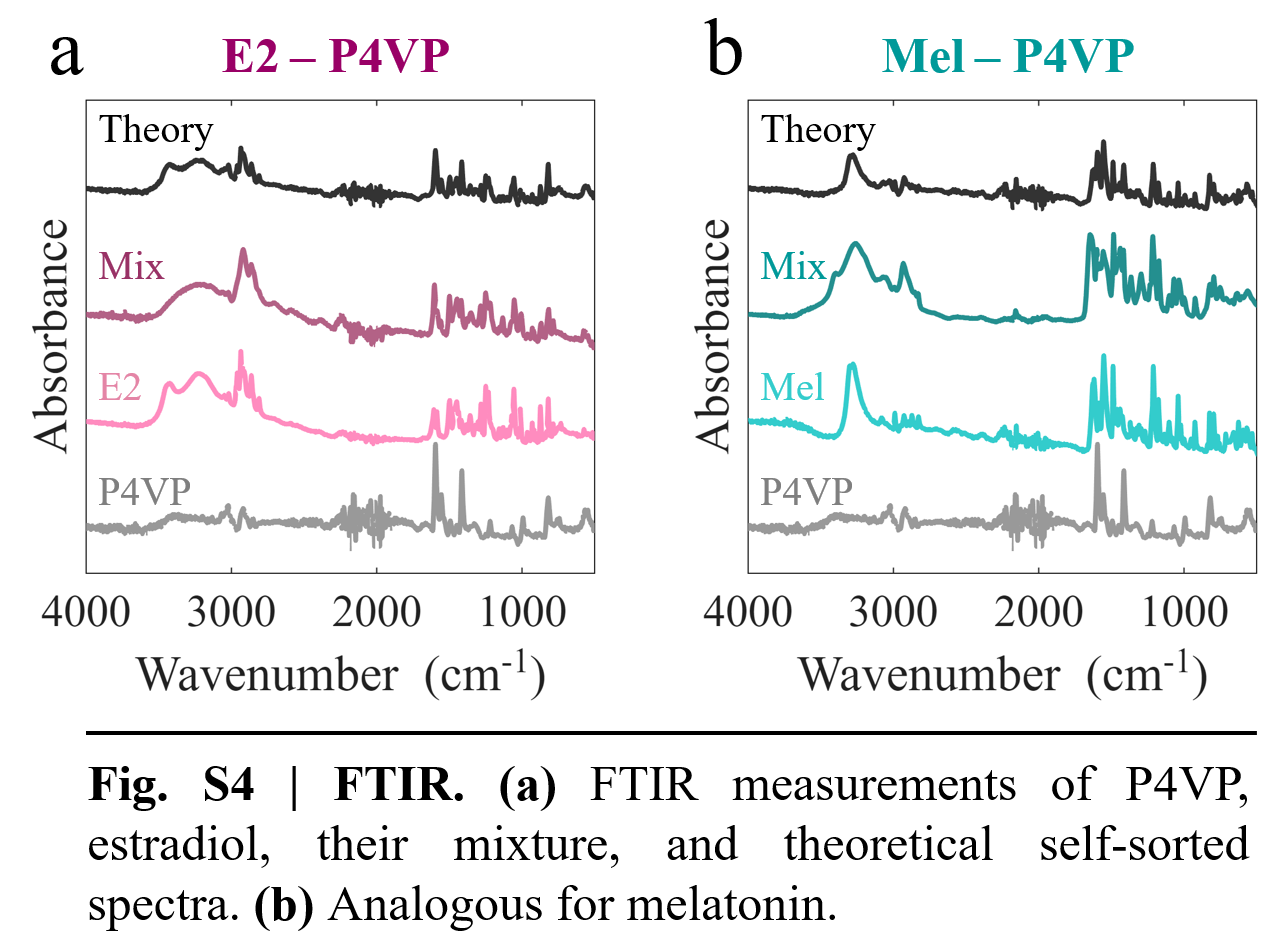

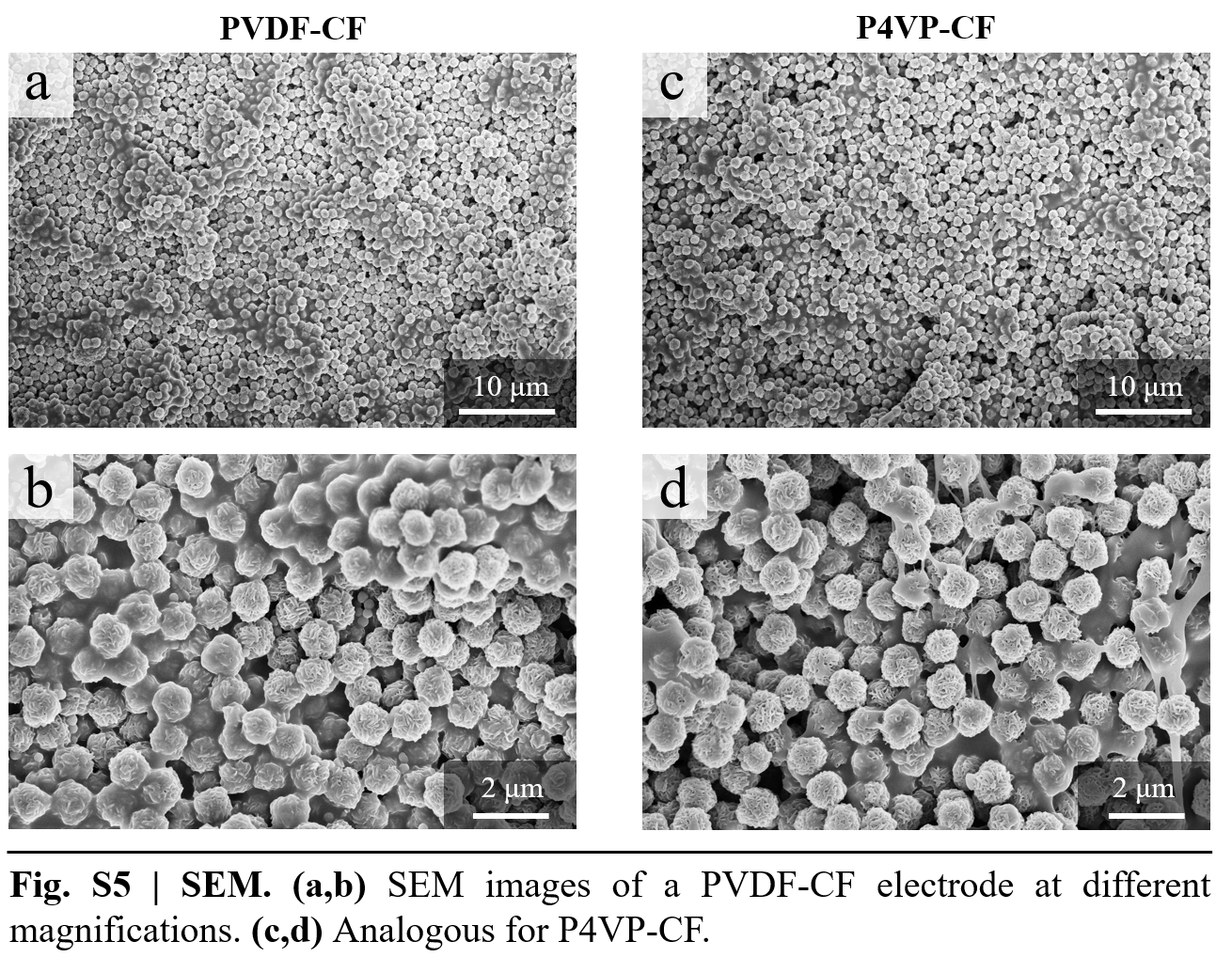

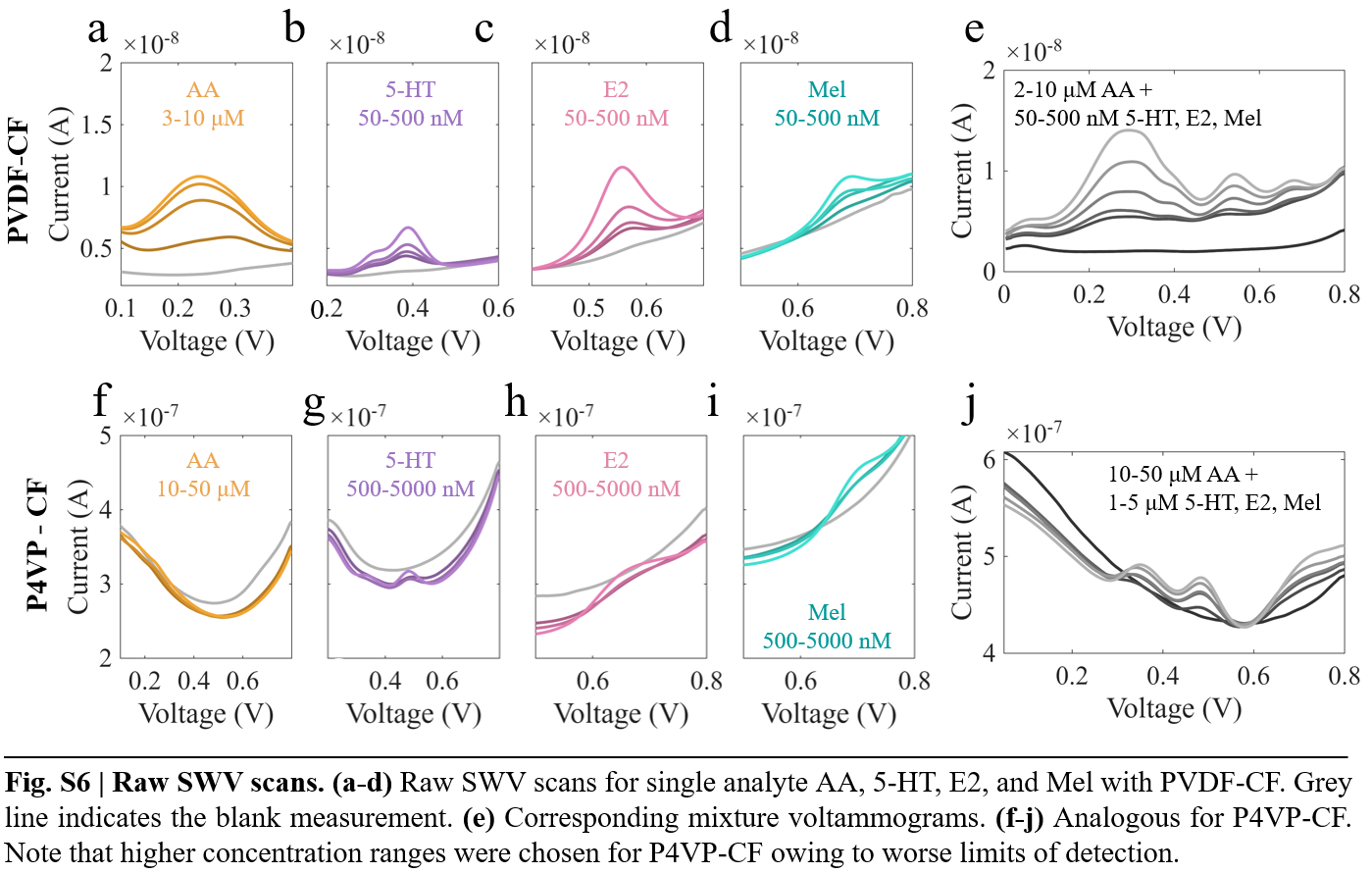
